# Deep Learning-Based High-Throughput Phenotyping Of Maize (*Zea mays* L.) Tasseling From Uas Imagery Across Environments

**DOI:** 10.1101/2024.06.24.600506

**Authors:** Nicholas R. Shepard, Aaron J. DeSalvio, Mustafa Arik, Alper Adak, Seth C. Murray, Jose Ignacio Varela, Natalia de Leon

## Abstract

Flowering time is a critical phenological trait in maize (*Zea mays* L.) breeding programs. Traditional measurements for assessing flowering time involve semi-subjective and labor-intensive manual observation, limiting the scale and efficiency of genetics and breeding improvement. Leveraging unoccupied aerial system (UAS, also known as UAVs or drones) technology coupled with convolutional neural networks (CNNs) presents a promising approach for high-throughput phenotyping of tasseling in maize. Most CNN image analysis is overly complicated for simple tasks relevant to plant scientists. Here a methodology for extracting tasseling from RGB imagery using a CNN-based approach was applied to 220 hybrids and 30 test lines grown in eight diverse environments (Wisconsin and Texas, U.S.A.) then validated through an unrelated set of hybrids. Overall accuracies of .946, .911, .985, and .988 were obtained for classifying maize images with or without tassels from College Station, TX in 2020; College Station, TX in 2021; Arlington, WI in 2021; and Madison, WI in 2021 respectively. By employing deep learning techniques, larger volumes of phenotypic data can be processed enabling high-throughput phenotyping in breeding programs. Although large datasets are required to train CNN models, the proposed methodology prioritizes simplicity in computational architecture while maintaining effectiveness in identifying flowered maize across diverse genotypes and environments.

## 1 Introduction

Achieving crop improvement, including higher yields, hinges on effectively measuring and managing various phenotypic traits that collectively contribute to yield, quality, and profitability. Field-based manual phenotyping faces scalability and time constraints, especially when traits of interest are only detectable during critical biological windows (Aasen et al. 2020). Modern breeding practices integrate high throughput phenotyping and genotypic data to establish comprehensive genome and phenome-based analyses (Araus & Cairns, 2014, Rincent et al. 2018, Zhu et al. 2021). While genomic automation advancements have accelerated beyond Moore’s law (Poland and Rife 2012, Delseny et al 2010), two major automation bottlenecks persist in phenomics: extracting phenomic measurements and phenotypes from images and efficiently analyzing the vast volume of phenotypic data acquired (Furbank & Tester 2011, Minervini et al. 2015, Song et al. 2021). Detecting phenotypic variation in near-real-time and at scale poses challenges without automated approaches, given the time-consuming and potentially error-prone nature of manual methods. Various methods have attempted to address these challenges but often do not meet the levels of throughput required for large plant breeding programs and commercial breeding operations (Lu et al. 2017, Shao et al. 2021, A & Sangeetha, 2021, Murcia et al. 2021).

Maize (*Zea mays* L.) is a monoecious plant with tassels containing the male anthers, which produce pollen, at the apex. Maize exhibits considerable variation in flowering time among both inbred lines (Buckler et al. 2009) and hybrids across environments (Rattalino et al. 2011), resulting in germplasm with variation in flowering initiation-related traits such as the duration of the grain-fill period (Daynard et al. 1971). Female flowering (i.e. silking) is highly correlated with male flowering (Izzam et al. 2017). Flowering time (both female and male) serves as a crucial indicator of the transition from vegetative to reproductive growth, with synchronous flowering in hybrid production fields essential for maximizing yield (Worku et al. 2016, Cárcova et al. 2000, Baum et al. 2019). Ensuring appropriate flowering time for a variety within its environment of production is also critical for optimizing hybrid seed production using inbred lines, which can be difficult where photoperiod associated variation occurs (Adak et al. 2021b). Conventional manual scoring of male flowering requires physically walking the field every day or every few days (with revisit time directly impacting accuracy) and manually estimating when 50% of plants in a plot show anthers (anthesis) or silks (silking) (Andersen et al. 2005, Mace et al. 2013, Khan et al. 2022). Transitioning from manual scoring of flowering in populations to measurement by unoccupied aerial system (UAS, also known as UAVs or drones) flights could dramatically reduce labor and time. UAS platforms can readily capture high-quality images, including tassel initiation, facilitating automated detection of maize tasseling in breeding fields (Kurtulmuş & Kavdir, 2014, Lu et al. 2016, Karami et al. 2021). While tassel initiation and anthesis are not identical, they are highly correlated, especially in material with large flowering windows (Warrington & Kanemasu, 1983), such as in exotic introgression and pre-breeding programs.

Incorporating artificial intelligence (AI) approaches, specifically deep learning (DL) applications, in plant breeding presents a promising approach to address the volume of unanalyzed or uninterpreted phenotypic data (Ubbens & Stavness, 2017, Namin et al. 2018). Previous DL applications in automated phenotyping have primarily been limited to controlled environment or ground-based field image acquisitions, which are often stationary, manually operated, or otherwise impractical for adoption at scales needed in genetic and breeding studies (Ye et al. 2012 Mirnezami et al. 2021, Shao et al. 2023). Studies employing deep learning techniques have shown promise in automating phenotyping tasks such as maize tassel morphology and development (Yu et al. 2022, Zhang et al. 2023), tassel counts (Lu et al. 2017, Lu et al. 2020, Zan et al. 2020), and the effect of tassels on leaf area index estimation using vegetation indices (Shao et al. 2023). However, these methods are constrained by either their destructive or non-scalable methodology or datasets to the thousands of diverse plots screened in field breeding and genetics programs (Alzadjali et al. 2021).

Recent advancements in deep learning have expanded the scope of automated phenotyping by applying modern techniques to existing datasets. These techniques include the use of k-means algorithms, which are unsupervised learning methods for clustering data into K groups based on similarity (Kumar et al. 2021). Similarly, K-nearest neighbor (KNN) algorithms, which classify objects based on their similarity to neighboring objects, have also been utilized in automated phenotyping tasks (Fitria Widiawati et al. 2018). Furthermore, convolutional neural networks (CNN) have emerged as powerful tools in automated phenotyping due to their ability to learn features directly from raw data, such as images (LeCun et al. 2015). Variants of CNNs, including Fast R-CNN and Faster R-CNN, have been explored for diverse object detection and classification tasks including plant phenotyping (Ren et al. 2015, Lu and Cao, 2020, Liu et al. 2020). While Faster R-CNN offers improved performance, it also requires higher computation which is a barrier.

When considering the choice of utilizing a pre-trained model for phenotypic analysis, the volume of datasets available in an immediately usable state do not necessarily resemble the data formats generated internally within different research programs. Unlike some pre-trained models, which exhibit a relatively high level of complexity and require a steep learning curve, alternative approaches and models can offer a more approachable and modifiable framework. This simplicity reduces the barrier to understanding but also provides improved interpretability mitigating the ’black box’ effect associated with more complex deep learning models. Moreover, employing a simpler model allows for in-house modifications tailored to specific research needs.

While advanced supervised models such as Faster R-CNN may offer sophisticated features, the additional computational requirements do not necessarily yield significant improvements in results compared to simpler alternatives (Rodene et al. 2024). Many cloud-based data services charge for the management of supervision during the learning process, adding to overall cost and complexity. Basic CNN architectures implemented using frameworks like Keras provide a more accessible, less computationally intensive, and easily applicable solution. The Keras CNN architecture used here was believed to be sufficient for the task while offering scalability and transferability across RGB images that differ in natural lighting conditions, resolution and environment. We hypothesize that deep-learning approaches for high-throughput tassel phenotyping are achievable without requiring highly-complex computational architecture which can lead to easier incorporation and scalability across years and locations.

## 2 Materials and Methods

### 2.1 Genomes to Fields Initiative (G2F) Maize Dataset

#### 2.1.1 Field Design

Images from field experiments were collected in three fields in College Station, TX, one field in Arlington, WI (WIH2) and one field in Madison, WI (WIH1) as part of the Genomes to Fields Initiative (G2F) in 2020 and 2021. Additional information on this experiment can be found in Lima et al. (2023). In brief, 220 hybrids and 30 test lines were grown in a randomized complete block design (RCBD) with two replicates, each plot comprising one hybrid with two rows (500 plots per environment). In College Station, TX and in Arlington, WI the hybrids were all crossed with tester parent PHZ51. In Madison, WI the hybrids grown had been crossed with tester parent PHP02. The specific data set in College Station, TX fields are as described in Adak et al. (2021a). In Texas the 500-plot hybrid trial was separately evaluated under three separate management conditions for 1500 plots in total: TXH1 is defined as having optimal management and planting date, TXH2 is defined as dryland and reduced fertilizer with optimal planting date, and TXH3 is defined as optimal management but one month delayed planting date.

#### 2.1.2 Data Collection

UAS data acquisition of RGB images was made using a DJI Phantom 4 Pro V2.0 (SZ DJI Technology Co. Ltd. Shenzhen, China) equipped with a 1-inch CMOS RGB sensor with a mechanical shutter. Using the DJI GS Pro application, flight missions were planned with the following parameters: 25m elevation (above ground), 90% forward overlap, 80% side overlap, flight speed of 1.2m/s, and a shutter interval of 2.0s. Raw images were sorted into separate folders named according to each flight date in preparation for construction of orthomosaics within Agisoft Metashape (Agisoft LLC). Lower flight heights allow for higher ground sampling distance/resolution.

#### 2.1.3 Data Processing

A summary of orthomosaicking and georeferencing procedures is described as: a) folders containing RGB images were loaded into Metashape; b) photo alignment was conducted using 60,000 key points and 0 tie points with referenced preselection; c) an initial bundle adjustment was performed to optimize the f, cx/cy, k1, k2, k3, p1, and p2 distortion parameters of the lens; d) iterative model error reduction procedures (termed “gradual selection” in Metashape) were performed to remove erroneous points from the sparse cloud; e) ground control points (GCPs) were imported and manual alignment of their locations within at least six RGB images was performed followed by selecting the “update” option within Metashape to integrate the GCPs into the model; f) camera alignment was performed again with all available distortion parameters; g) the dense point cloud was processed with “moderate” depth filtering at “medium” quality with all other settings left as default; h) color calibration was performed using the sparse cloud as the source and the “calibrate white balance” option was also checked; i) the digital elevation map (DEM) was calculated using the dense cloud as the data source with all other settings left as default; j) the orthomosaic was produced using the DEM as the surface with all other settings left as default. All georeferenced final products were exported in World Geodetic System 1984 (WGS84) datum (specifically, UTM Zone 14N for Texas and 16N for Wisconsin) coordinates. Orthomosaics and DEMs were exported from Metashape and saved with the .tif extension, while the dense point cloud was saved with both .las and .laz file extensions.

In total, the dataset consisted of 15,000 .tif images collected from thirty flights conducted across multiple years and environments. Only flights during the initiation of tasseling were selected. In 2020, four flights of fields TXH1 & TXH2 and three flights of TXH3 were used (Figure 1a). In 2021, five flights of TXH1 & TXH2, three flights of TXH3, three flights of WIH1, and three flights of WIH2 were conducted. Each flight took a composite of RGB images containing all 500 plots at differing quality. Of the 15,000 total images, the dataset became 13,491 images after removing images that were greater than 50% soil or had no recorded DTA (Table 1).

**Figure 1:**
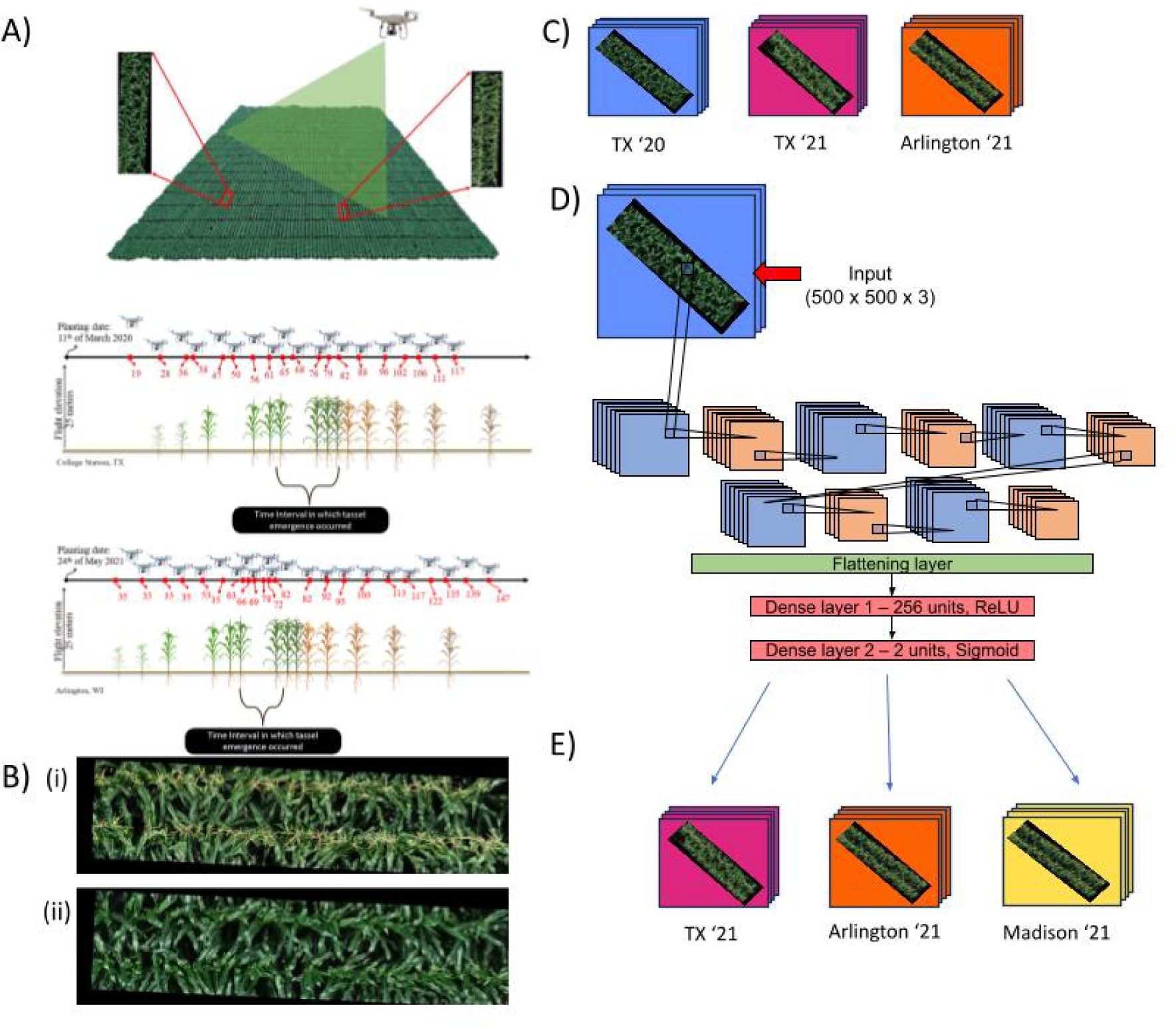
Summary of initial methods. **A)** UAS images were used to generate orthomosaics of the field across flight dates (as days after planting, (DAP)) for College Station, TX in 2020 and Arlington, WI in 2021 containing images of maize with and without tassels. The highlighted window contains the flowering window used. **B)** Row plots were then cropped into images, consisting of either a (i) flowered or (ii) unflowered plot. **C)** Images from College Station, Texas in 2020 and 2021 as well as Arlington, Wisconsin in 2021 were visually scored using a 0 or 1 for flowering status. **D)** Keras-based CNN model was trained iteratively on a randomly sorted 80% subsample of preprocessed images in the RGB spectra from only College Station, Texas plots in 2020 alone with dimensions 500 x 500 pixels x 3 (RGB). **E)** Model predicted classes for unseen images that were withheld from training from College Station, Texas in 2021; Arlington, Wisconsin in 2021; and Madison, Wisconsin in 2021.

**Table 1.**
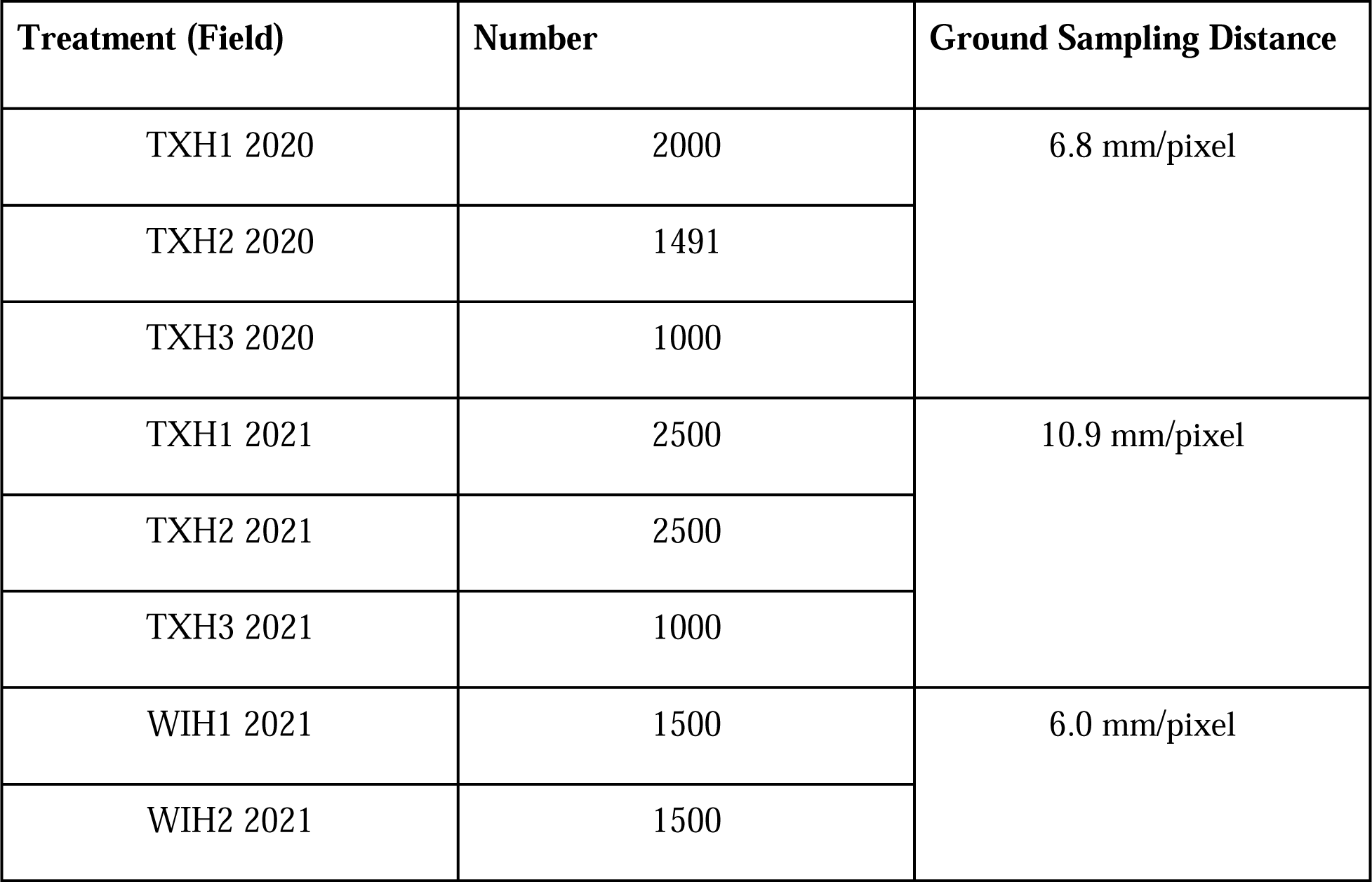
Image distribution and resolution.

Further processing of acquired images involved pre-processing steps using R and Python coding including resizing, removal of alpha channels, and normalization of pixel values (Supplemental data 1). Soil correction was performed using the “fieldimageR” package to remove soil artifacts and ensure accurate phenotypic analysis by only giving pixels containing maize plants value (Matias et al. 2020).

#### 2.1.4 Manual scoring

To establish ground truth for model training, all images were visually scored by the author for the presence of tassels (Figure 1b, 1c) by categorizing flowering status into either ‘0’ meaning less than fifty percent flowered or ‘1’ meaning more than fifty percent flowered, categories as estimated visually. The date of 50% anthesis, as days to anthesis (DTA) and 50% silking as days to silk (DTS), were recorded from visual observations manually walking the field every two to three days once any tassels were first observed. The tassel initiation date was unfortunately not recorded in these manual scorings but were expected to be a few days before DTA and highly correlated. Days to tasseling (DTT) was calculated as the days after planting (DAP) corresponding to the first flight date where tassels are visually present in more than 50% of maize plants.

#### 2.1.5 Summary Statistics

The correlation between DTT and DTA was determined in R. A random effects model ANOVA was performed using the lme4 package to determine repeatability measurements of DTA and DTT for all environments excluding Madison where a different tester was used according to Eq. 1 as well as by treatment according to Eq. 2 in Texas where there were multiple years and Eq. 3 in Wisconsin where there was only one year.

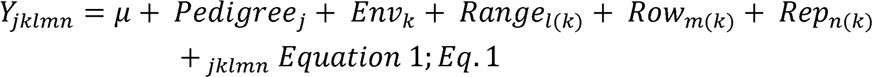

Y is a vector of length 230 (the sum of the number of hybrids) denoting each trait value (DTT or DTA) in DAP for each hybrid in one of eight environments. Here denotes the grand mean; Pedigree denotes the overall effect of the j^th^ maize hybrid, Env denotes the overall effect of the k^th^ treatment environment; Range denotes the effect of the l^th^ range nested in the k^th^ environment; Row denotes the effect of the m^th^ row nested in the k^th^ environment; Rep denotes the n^th^ replication nested in the k^th^ environment; and denotes the error term addressing non-explained variability.

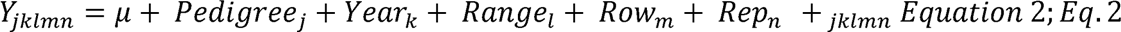

Equation 2 differs from Equation 1 such that Year is included as an effect while Range, Row, and Rep are no longer nested effects as analysis is being performed on a per environment basis.

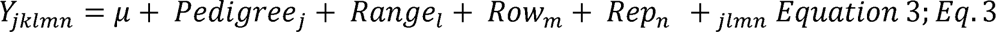

Equation 3 differs from Equation 1 such that each environment (year by location) is analyzed separately and therefore Range, Row, and Rep are no longer nested effects.

Repeatability was calculated according to Eq.4. Here, 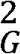 and ^2^ indicate the variance components of the effects of genotype and the error term, respectively, and n is the number of replicates:

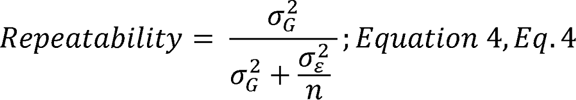

### 2.2 Flowered Plot Detection CNN

#### 2.2.1 Python Code

The essential components to recreating this analysis method are 1) the python code itself, 2) a folder containing logically named images from all flights, and 3) an array of values containing the corresponding flowering status label for each individual image file used in training. Spyder 5.4.3 was used via the Anaconda Navigator environment manager. The version of python running in Spyder was 3.11.5, with the following packages: pandas, numpy, os, keras, tensorflow, seaborn, matplotlib.pyplot, and imageio.v3. Image files were named with a date code format “YYYYMMDD-’’ which allowed for individual years and locations to be subdivided. Given large differences in flight dates between environments, specific regions could be selected by year (e.g. 2020 or 2021) or by the combination of year and month together (e.g. 202105 or 202107). Images were read, resized to 500 x 500 pixels, and checked to ensure they had the dimensions 500 x 500 x 3 representing only the RGB bands and removing any alpha channels before being stored in an array. This array had all pixel values of ‘NA’ set to ‘0’ (an artifact of removing soil with fieldImageR) before normalizing all pixel values from .tif format images by dividing by 255 (the maximum pixel brightness value) such that all pixel values fell within a range of [0, 1].

Each model was repeated ten times including the randomly iterated train/test split for image files from Texas in 2020 which was 80% training 20% test. Texas 2020 was the only set used for training. Training of the model occurred over fifty epochs, with subsequent classifications made on previously unseen images from Texas 2020, Texas 2021, and Wisconsin 2021 datasets. Performance of the DL approach was analyzed according to the following equations:

Precision (Eq. 5) measures the accuracy of positive predictions made by the model. It was calculated as the ratio of true positive predictions to the total number of positive predictions made by the model, including both correct and incorrect positive predictions.

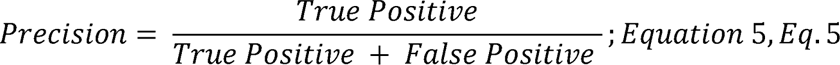

Recall (Eq. 6) measures the completeness of positive predictions made by the model. It was calculated as the ratio of true positive predictions to the total number of actual positive instances in the dataset. Mathematically, recall is represented as:

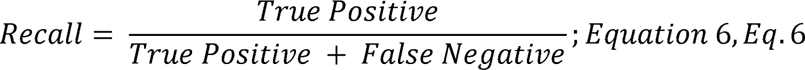

The F1-score (Eq. 7) is the harmonic mean of precision and recall. It provides a balance between precision and recall, giving equal weight to both measures. F1-score is calculated using the following formula:

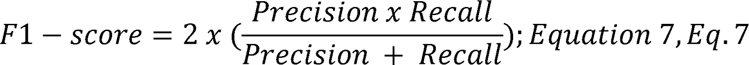

Accuracy (Eq. 8) measures the overall correctness of the model’s predictions. It was calculated as the ratio of correct predictions (both true positives and true negatives) to the total number of predictions. Mathematically, accuracy is represented as:

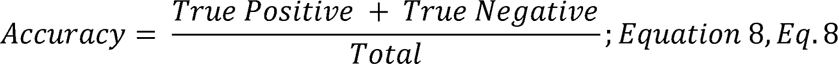

In order to properly save metrics, empty lists for evaluation, metrics, confusion matrices, predicted labels, and F1-scores were initiated. Flowering status values were one-hot encoded before the CNN model was executed. The CNN model architecture was constructed after performing hyperparameter optimization (Figure 2) and is as described in Table 2. Hyperparameter optimization was performed using the Optuna package in Python (Akiba et al. 2019) and determined that the most important hyperparameter was having five conv2D layers. The goal of hyperparameter optimization is to isolate the specific combination of adjustable variables that will produce the best accuracy while minimizing loss. Loss shows the difference between the predicted value and the ground truth value, and the difference between loss and validation loss is a useful metric for gauging over- or underfitting. We included the line ‘tf.keras.backend.clear_session()’ at the end of the loop in order to optimize memory usage.

**Figure 2:**
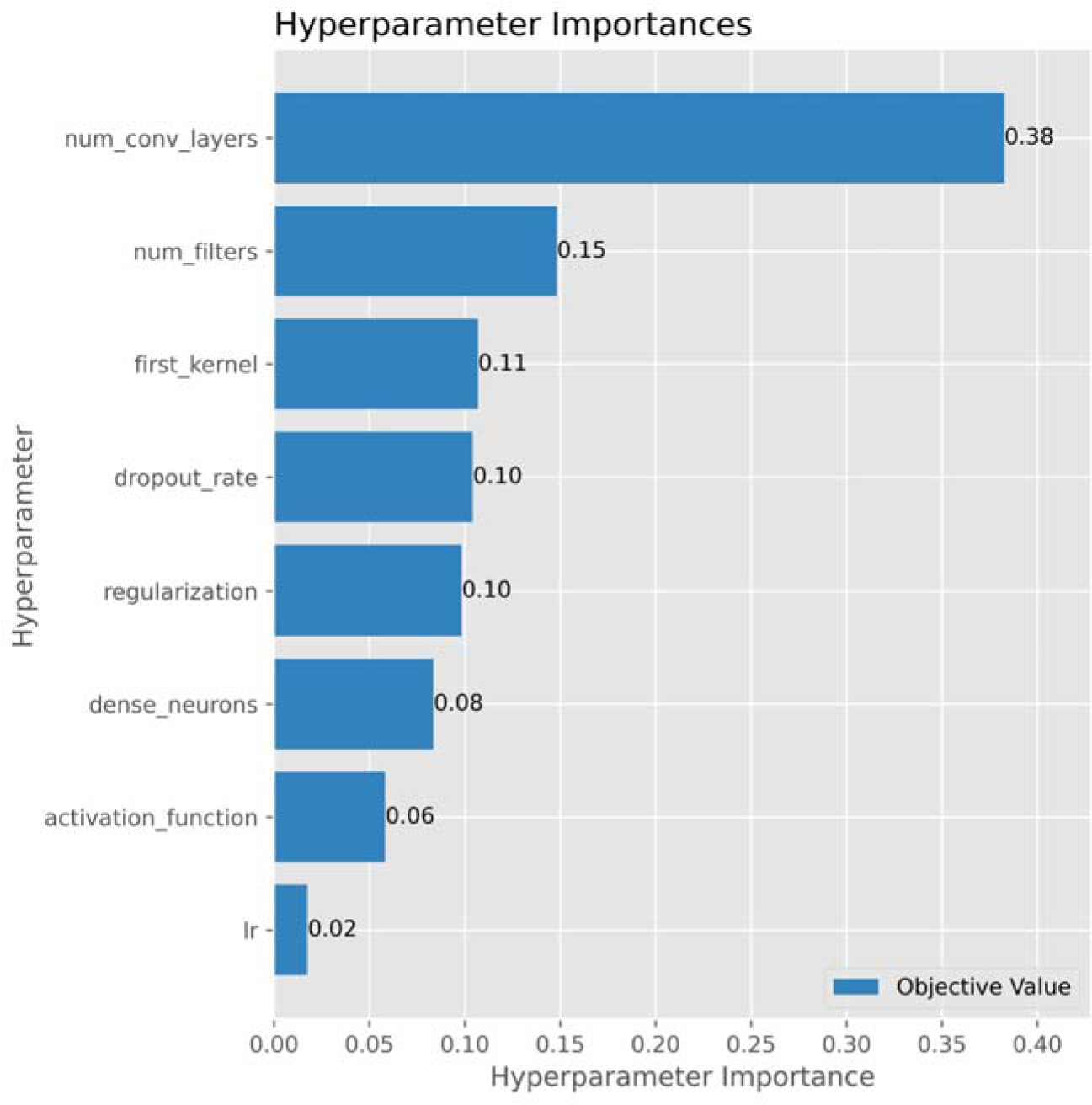
Hyperparameter importance as determined by Optuna. OptunaV3 was performed iteratively over fifty trials examining the importance of all listed tunable hyperparameters.

**Table 2.**
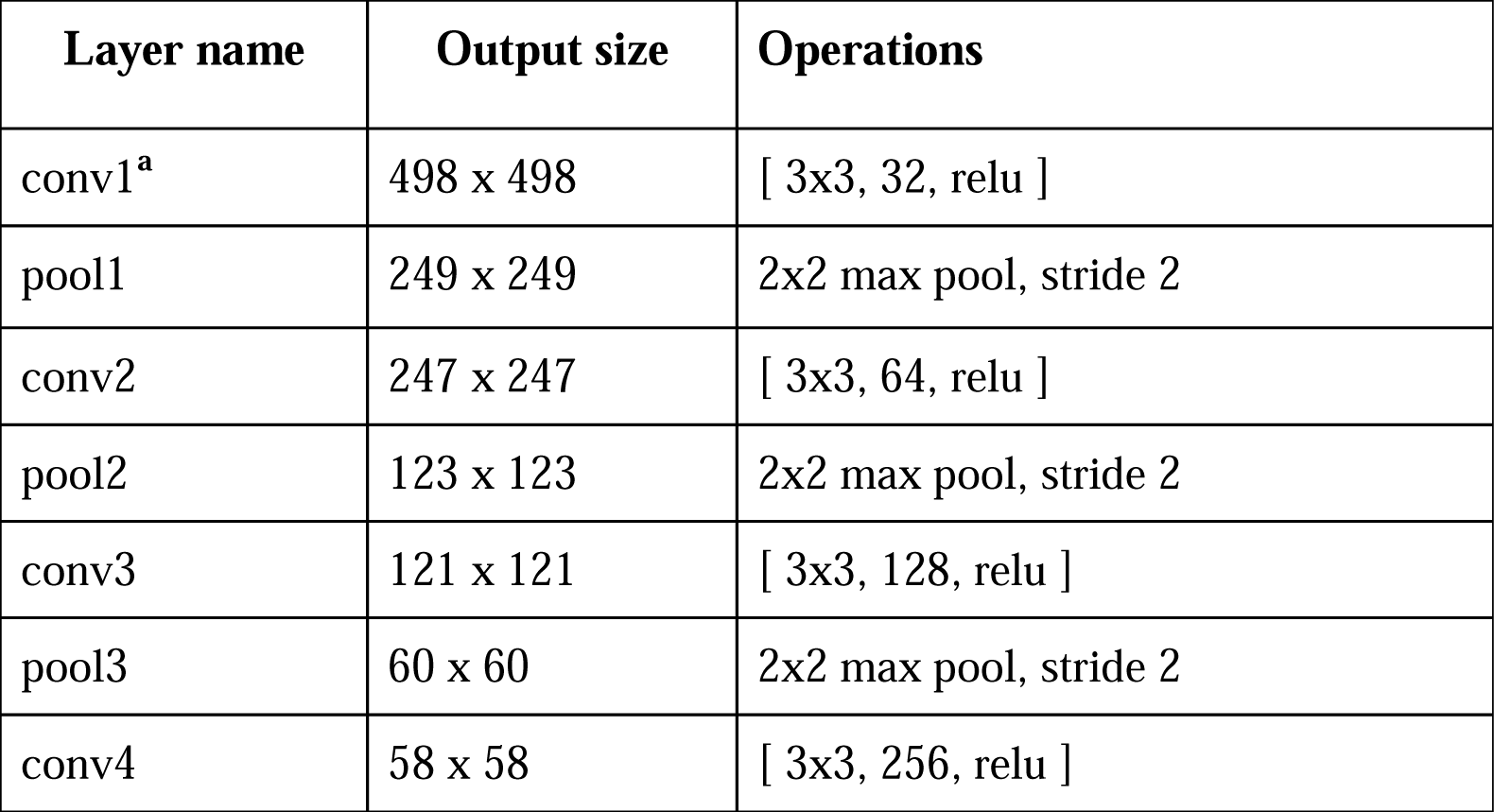
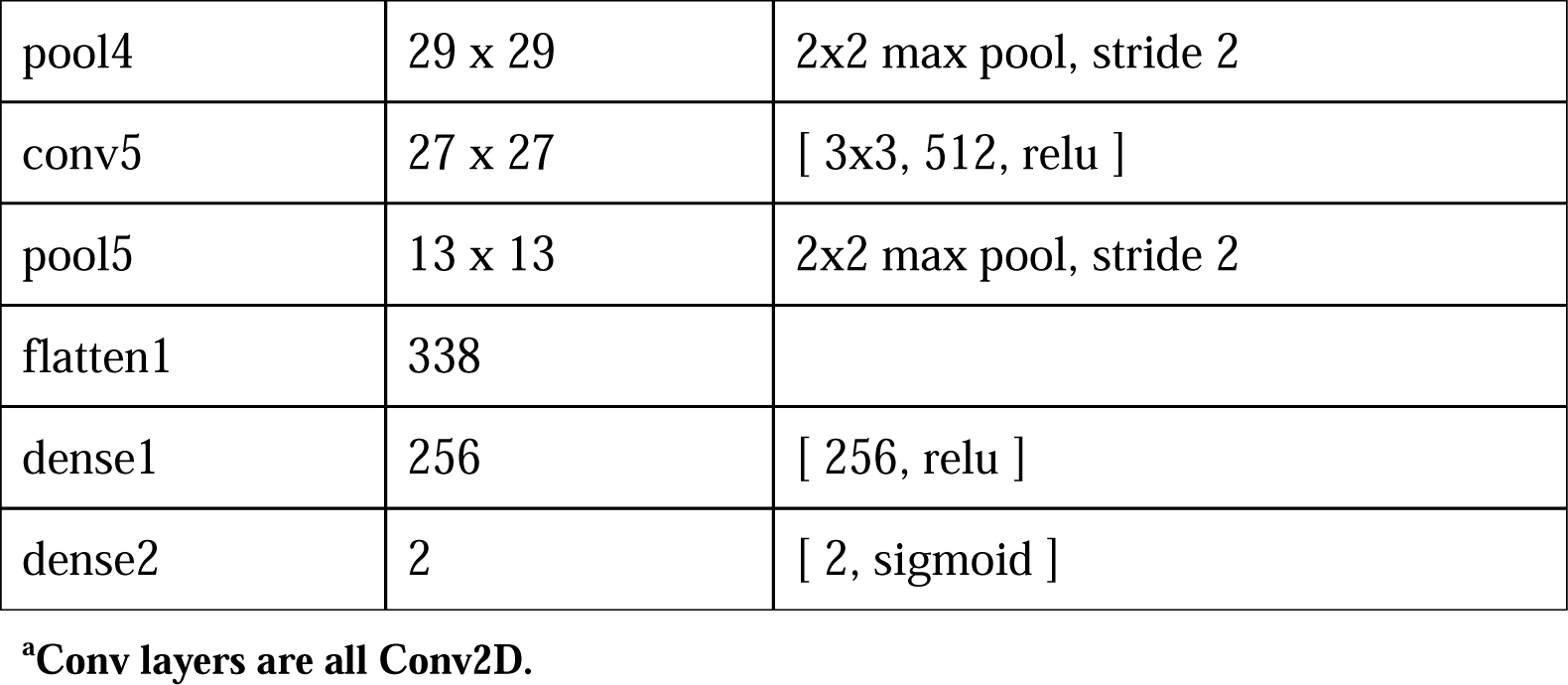
CNN Architecture.

Evaluation of model performance included the generation and storage of confusion matrices and saliency maps to visualize classification results. Saliency maps were created by copying the model to extract and visualize specific layers for specific images, allowing the export of layers that show activation for either maize showing tassels or not. Metrics were concatenated into structured lists and exported to CSV files for comprehensive analysis and documentation. F1-scores were then compared against previously published deep learning studies. Labels lists were generated and sorted to show which flight days were the first to record tasseling.

## 3 Results

### 3.1 Summary Statistics

#### 3.1.1 Statistical Analysis of DTT and DTA

Correlation (r^2^) between DTT and DTA based on the data in Lima et al. (2023) was found to be 0.71 (Figure 3). ANOVA was performed on trials all using the same genotypes (TXH1, TXH2, TXH3, and WIH2 datasets) for DTT and DTA (Table 3a), as well as for WIH1 individually (Table 3b), and all environments individually (Supplementary Tables 1-10).

**Figure 3:**
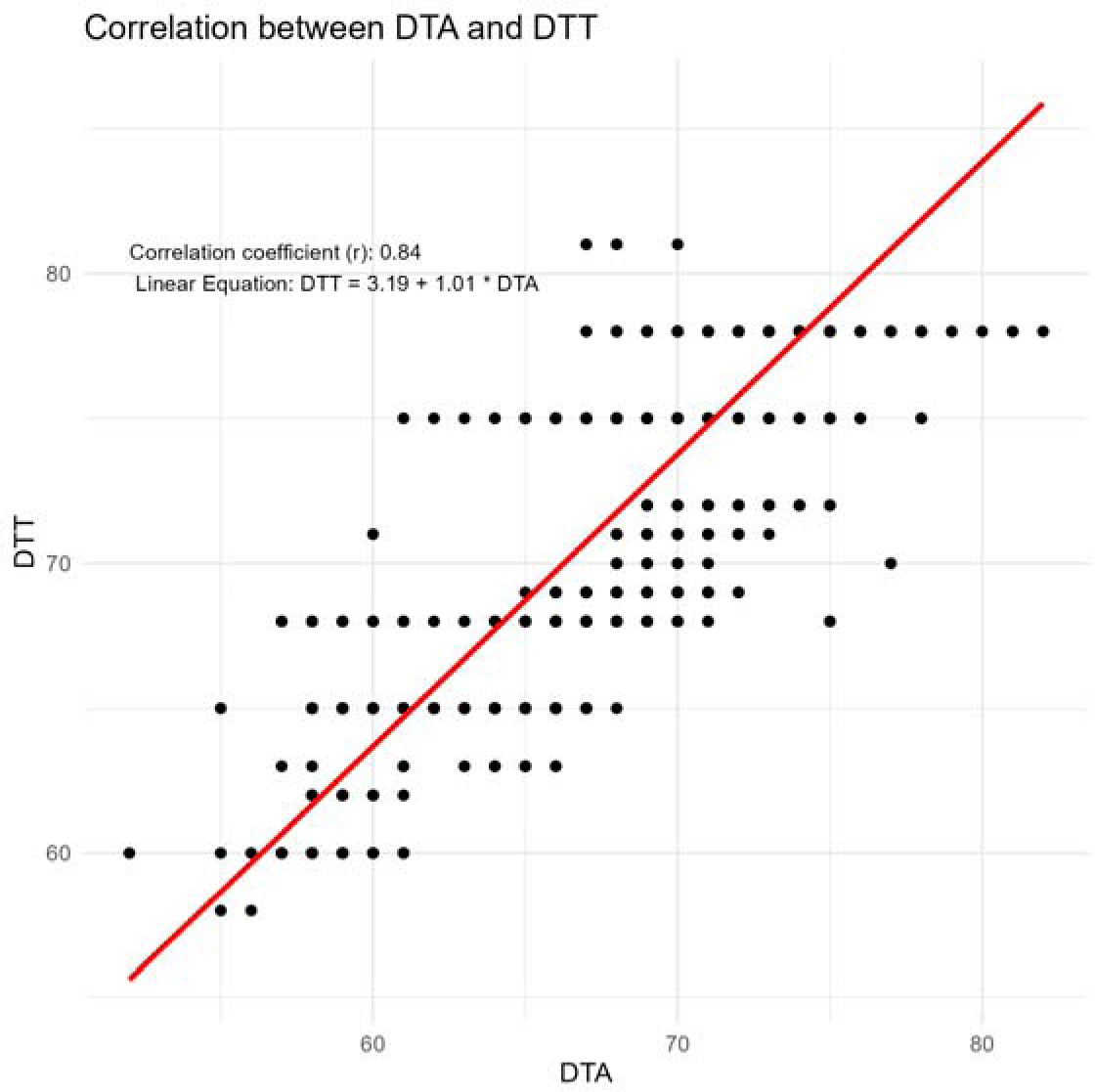
Correlation between DTT and DTA in both WI and TX in 2020 & 2021. Correlation between days to tasseling (DTT) and days to anthesis (DTA).

**Table 3a.** ANOVA results for DTA and DTT of the entire dataset excluding Madison.

|  | DTA<br>vcov | DTA<br>sdcov | DTA Percent | DTT vcov | DTT sdcov | DTT Percent |
| --- | --- | --- | --- | --- | --- | --- |
| Pedigree | .632 | .957 | .028 | .413 | .643 | .012 |
| Treatment:Row | <.001 | <.001 | <.001 | <.001 | <.001 | <.001 |
| Treatment:Range | .092 | .302 | .004 | .251 | .501 | .008 |
| Treatment:Rep | .650 | .806 | .028 | .281 | .531 | .008 |
| Treatment | 16.24 | 4.03 | .705 | 26.68 | 5.16 | .802 |
| Residual | 5.42 | 2.33 | .235 | 5.62 | 2.37 | .169 |
|  | RMSE | RSquared | Repeatability | RMSE | RSquared | Repeatability |
|  | 2.247 | .764 | .651 | 2.299 | .831 | .541 |

**Table 3b.** ANOVA results for DTA and DTT for Madison, WI (WIH1).

|  | DTA<br>vcov | DTA<br>sdcov | DTA Percent | DTT vcov | DTT sdcov | DTT Percent |
| --- | --- | --- | --- | --- | --- | --- |
| Pedigree | .441 | .664 | .191 | .252 | .502 | .370 |
| Row | .276 | .525 | .119 | .053 | .231 | .078 |
| Range | <.001 | <.001 | <.001 | <.001 | <.001 | <.001 |
| Rep | .056 | .237 | .024 | <.001 | <.001 | <.001 |
| Residual | 1.53 | 1.24 | .665 | .377 | .614 | .552 |
|  | RMSE | RSquared | Repeatability | RMSE | RSquared | Repeatability |
|  | 1.08 | .335 | .365 | .480 | .448 | .572 |

### 3.2 CNN Results

#### 3.2.1 CNN Performance Evaluation

Overall the model was highly accurate (.948) at detecting tassels within the Texas 2020 dataset (Figure 4 A&B). It was also highly accurate (.911 and .981) on independent test data sets in Texas 2021 and Arlington, Wisconsin (Figure 4C). Precision, Recall, and F1-scores for corresponding datasets are presented in Figure 4D.

**Figure 4:**
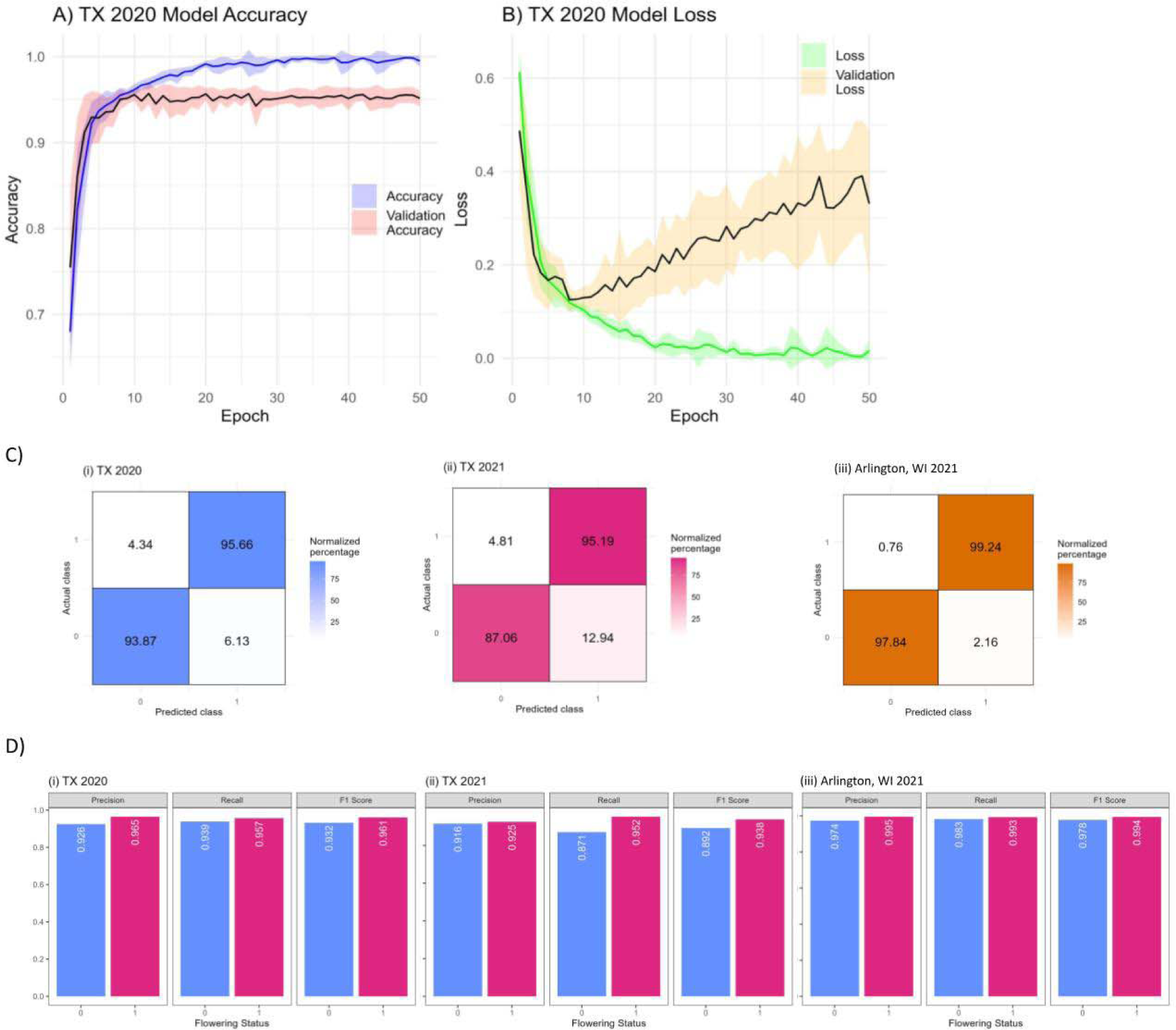
Model training & validation. **A)** Accuracy (blue) and validation accuracy (red) for the TX 2020 model which was trained on 80% of the images from College Station, TX in 2020. **B)** Loss (green) and validation loss (orange) for the TX 2020 model. **C)** Confusion matrices for unseen images in **(i)** TX 2020, **(ii)** TX 2021 and **(iii)** Arlington, WI 2021 all classified using the TX 2020 model. Rows of each confusion matrix are normalized such that each row sums to 100%. **D)** Precision, Recall, and F1-scores for each environment (**(i)** TX 2020, **(ii)** TX 2021 and **(iii)** Arlington, WI 2021) based on flowering status.

Saliency maps were generated showing the activation of different convolutional or dense layers for maize plants with and without tassels (Figure 5). Layers with tassels present show more activation than the same layer without tassels.

**Figure 5:**
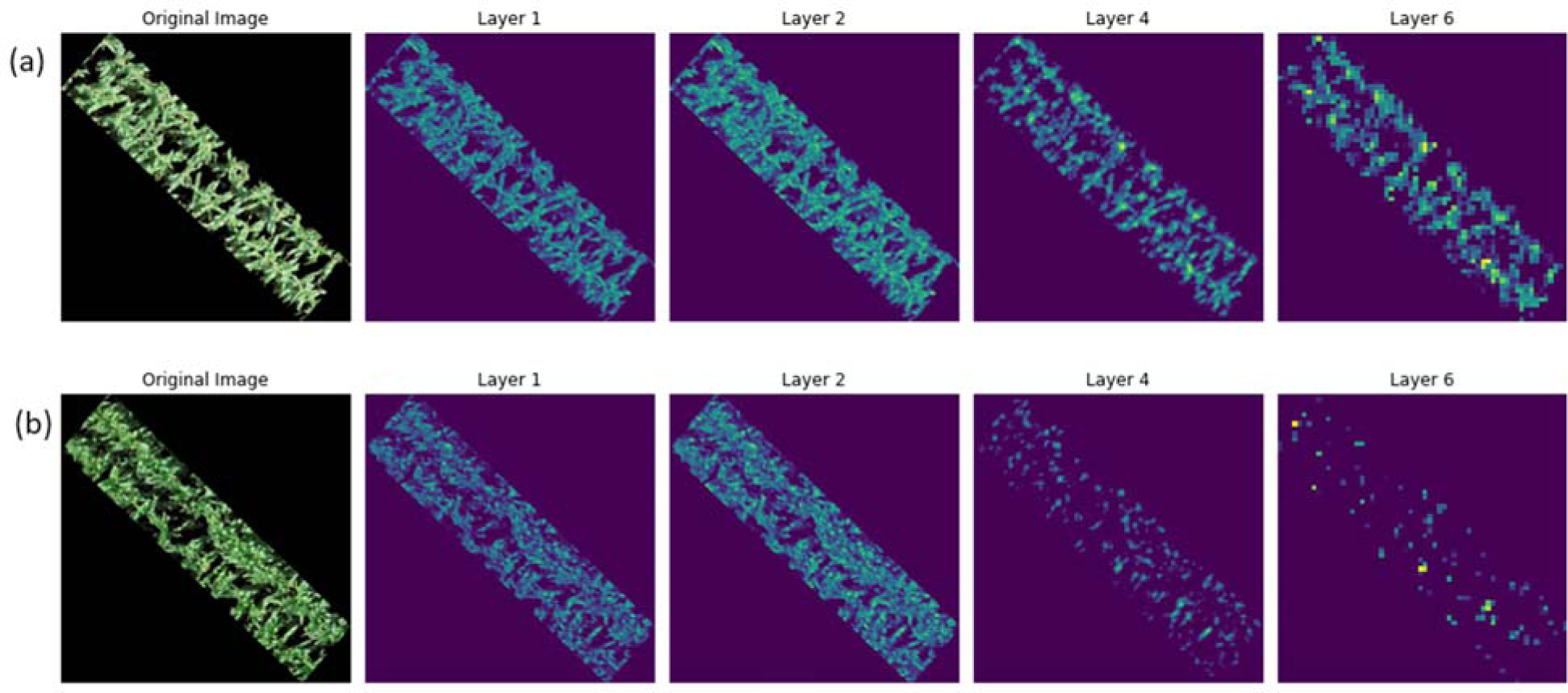
Saliency maps. Activation maps of a **(a)** flowered/with tassels and **(b)** non-flowered/without tassels in the cropped images

#### 3.2.2 Madison, WI Tassel Detection

Prediction of classes were generated for images of hybrids grown in Madison, Wisconsin in 2021 as part of G2F but were not previously used for training in our study. These used th same inbred lines but a different tester (PHP02) to create the hybrids. Images were scored manually and then labeled using ten replications of randomly partitioned train/test splits from the TX 2020 trained model. The model was very accurate (.988) at determining flowering status for the unrelated hybrids in Madison, Wi (Figure 6).

**Figure 6:**
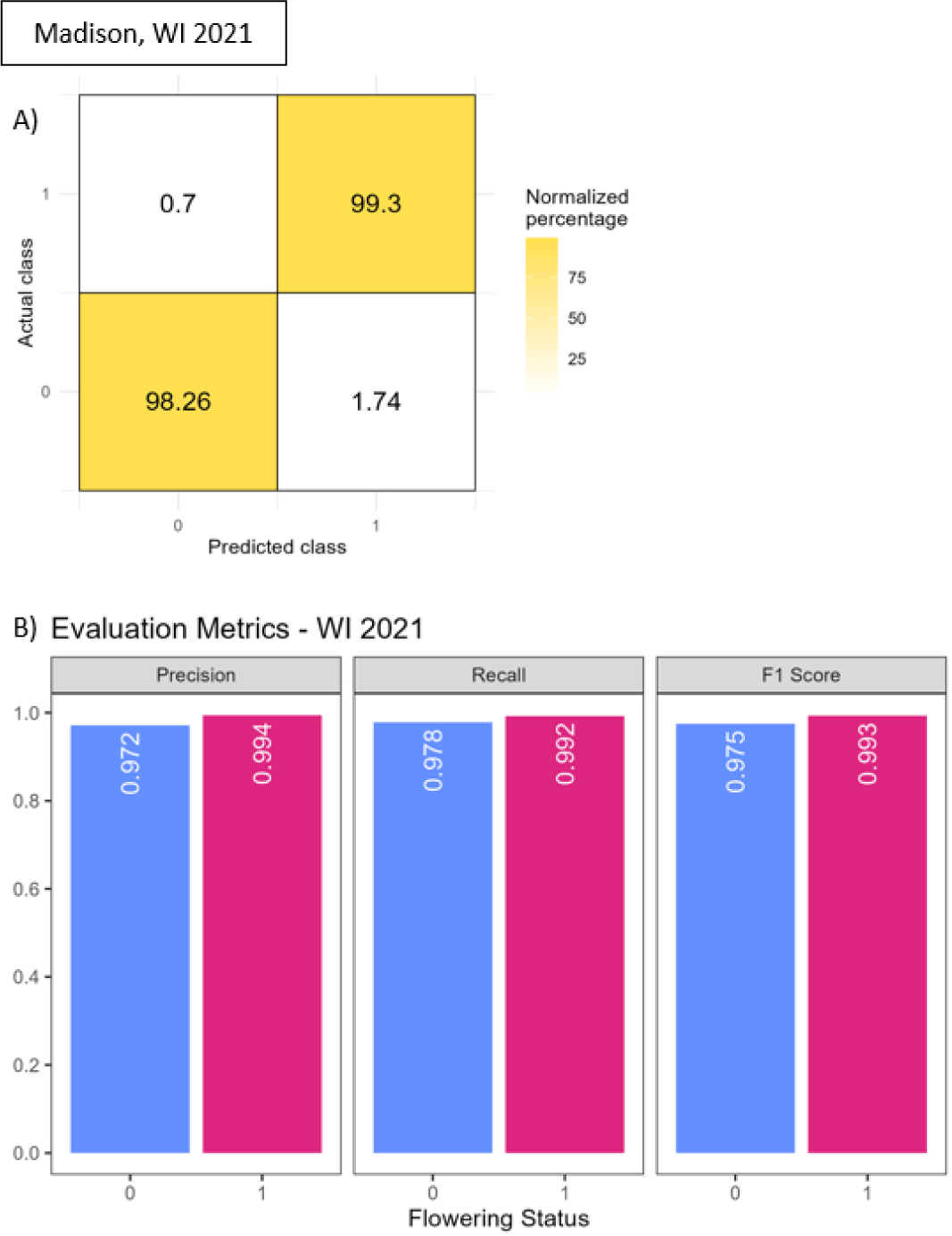
Madison, WI metrics. Deep learning metrics for the distinct hybrids in the Madison environment that were not present in College Station, TX or Arlington, WI. **A)** Confusion matrix for the actual and predicted classes of maize images from Madison, WI. **B)** Precision, recall, and F1-scores for maize images from Madison, WI based on actual flowering status.

## 4 Discussion

The CNN model used here, when trained on large datasets, demonstrated effectiveness in automating phenotyping tasks including the detection of tasseling plots in maize fields without necessitating more complex computational tools (Table 4). By automating the identification of flowering maize using UAS-based imagery, the objective was to replace the time-consuming and subjective process of manual recording days to anthesis or tassel identification from images with a more efficient and objective method. The limited image resolution of most UAS data sets, including this one, is insufficient to observe the presence of pollen released from anthers (DTA). However tassels are reasonable proxies and could be observed in these images.

**Table 4.** F1 scores of varying DL approaches that identify maize tassels. Higher (0-1) scores are better.

| Study | DL approach/architecture | F1-scores |
| --- | --- | --- |
| Zan, 2020 | RandomForest & VGG16 | .94 |
| Karami, 2021 | CenterNet, TSD, DetectoRS | .932, .899, .903 |
| Alzadjali, 2021 | Faster R-CNN (ResNet50) | .979 |
| Alzadjali, 2021 | TD-CNN (Inception v3) | .959 |
| Shepard, 2024 | CNN (Keras) | .96, .935, .99 |

**Table 5.**
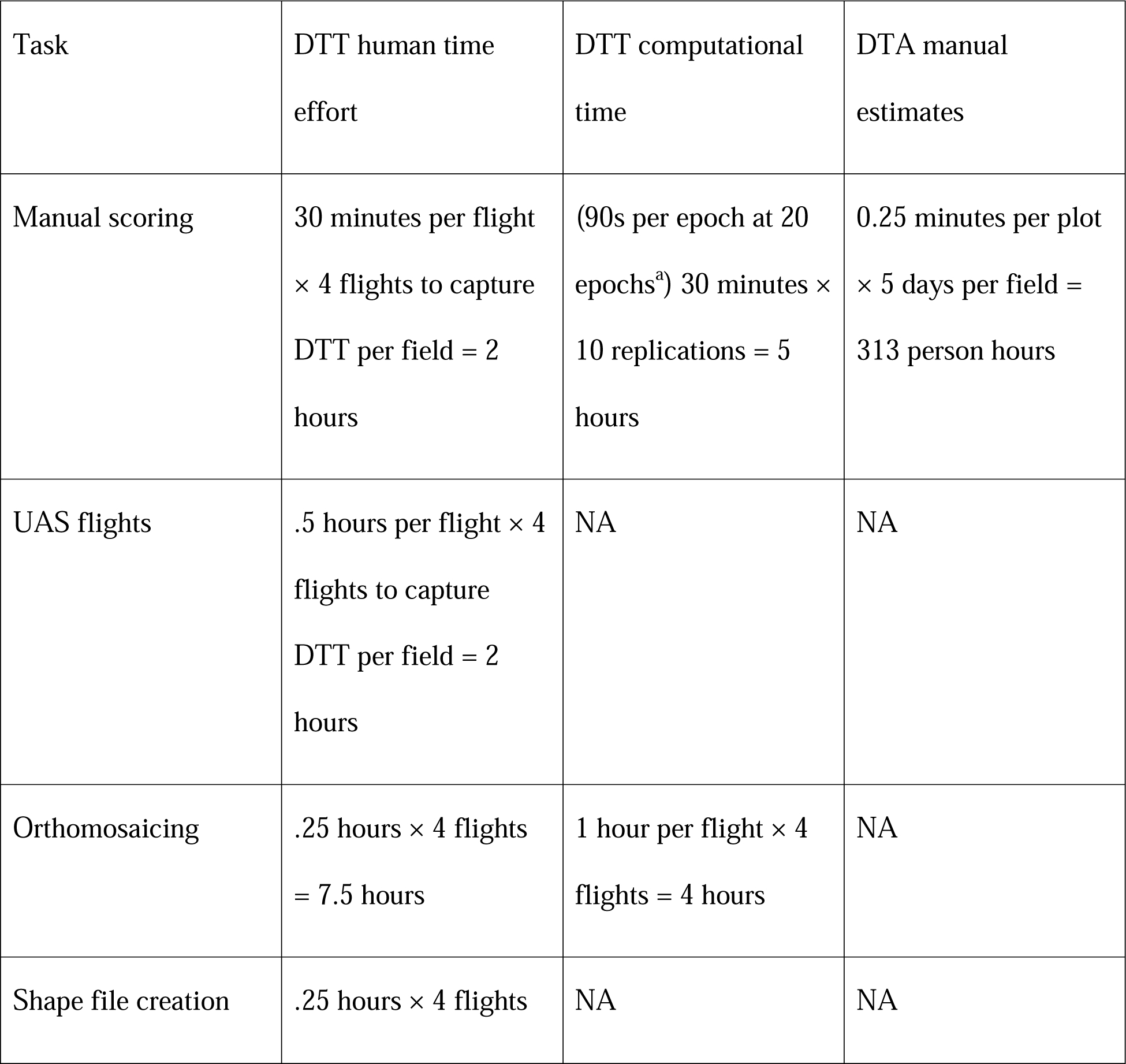
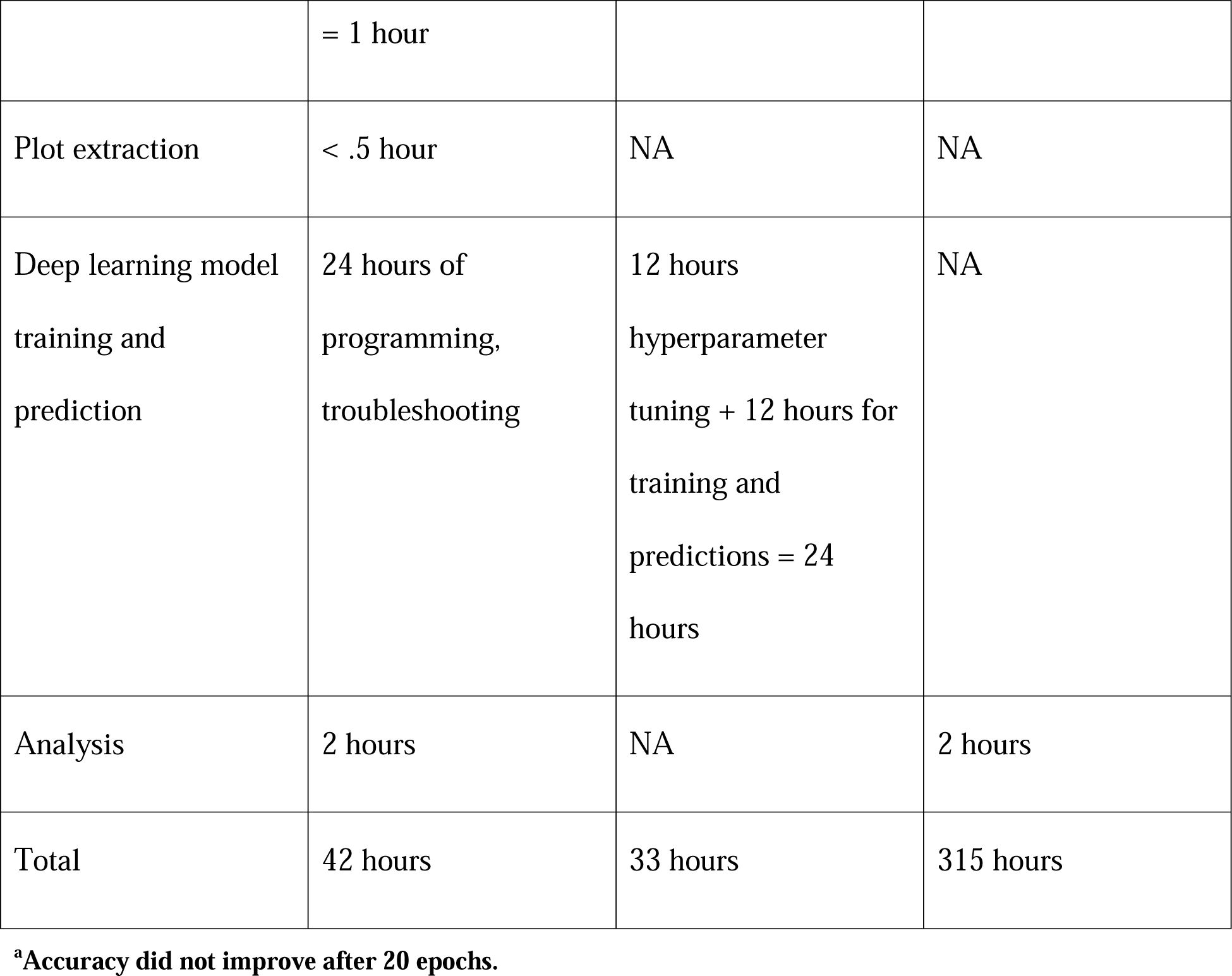
Time estimates for DTT by human and computer, DTA by human.

Beyond treating the image collection dates as independent, the date of tassel appearance could be extracted from these deep learning methods by arranging predicted classes by date and finding the first occurrence of a label indicating the detection of tassels, however this is only practical if the flights are spaced closely enough. Tassel emergence is a relatively rapid process, therefore in order to determine tassel date by UAS imagery flights should occur daily within the flowering window of all tested varieties. For example, if tassel initiation occurred at 73 days after planting but flights only occurred on days 72 and 78, the tassel emergence wouldn’t be captured until day 78. Conversely, if flights were made multiple times a day, the precision of tassel emergence to time within a day might be able to be detected. This would have relevance to quantitative genetic studies to dissect loci with smaller effects than a single day as it appears the majority of segregating flowering loci demonstrate in maize (Buckler et al. 2009). Alternatively, to get timing of tassel appearance with sparse UAS data, future deep learning models could go beyond classification to develop quantitative models for estimates of tasseling; either as the percentage of plants per plot tasseling or the percentage of the tassels emerged within plants in the plot. However, these approaches would require manual estimates on individual plants or manual quantitative estimates of how much the tassel has emerged to sufficiently train the model. Given the high repeatability observed for DTT (and DTA) demonstrated here, the manual and predicted classification of DTT was a high quality measure and it would make sense to invest in quantitative estimation.

Comparing resource investments between DTT and DTA is complicated, depending on technology available and used. The rough estimates here (Table 4) show that DTT took more effort, and higher education levels than DTA, which can easily be measured by student workers with limited education and training. However, it is worth noting that UAS collection and analysis technologies will continue to improve in speed and accuracy for DTT, while DTA can not be further scaled or improved. More importantly many other phenotypes and predictions can simultaneously be made using the same UAS data (Gano et al. 2024), such as plant height (Anderson et al. 2019, Pugh et al. 2018, Tirado et al. 2020), yield predictions (Kumar et al. 2023, Sunoj et al. 2021, Barzin et al. 2020), disease (Wu et al. 2019, Chivasa et al. 2021, De Salvio et al. 2022), and further enhance our understanding of the phenome (Murray et al. 2023). Nevertheless, a recent survey identified the “high cost of instruments/devices or software” and the “Lack of knowledge or trained personnel to analyze data” as important barriers to UAS adoption (Lachowiec et al. 2024).

The decision here to use a simple, modifiable model as opposed to pre-trained alternatives or one with higher computational requirements aligns with the goal of making deep learning-based phenotypic analysis more accessible and cost-effective without sacrificing accuracy. The affordability and versatility of RGB cameras further support this experience, enabling programs with limited resources to still benefit from incorporating more phenotypic data into research endeavors. Unlike hyperspectral cameras, which are more expensive and require specialized equipment, RGB cameras offer an opportunity to readily incorporate higher image resolution and more phenotypic data suitable for deep learning techniques. Programs using RGB can therefore capitalize on the potential for larger datasets, even with constrained resources, by implementing these methods more quickly. Multispectral camera data, incorporating a few additional bands over RGB with comparable speed and resolution, could further improve results and are worth investigating for DTT. However, this necessitates the use of more expensive technology, potentially prohibitive for smaller-scale programs and producers (Supplementary Table 11).

While coding this approach in Python, different situations were encountered where the architecture of the CNN would lead to erroneous predictions. It is important to regularly inspect activation maps for proper convolutional layer activation at the trait of interest as well as confusion matrices with predicted labels. In situations where the flowering status of unscored images is being classified, it is important to manually assess any image that does not receive a consistent predicted label. In doing this it was observed that orthomosaic stitching created artifacts in certain segments of the field, generally due to windy conditions or poor overlap between images. Furthermore, poor image resolution led to less consistent classifications.

The development of a user-friendly methodology suitable for deployment among a diverse range of users or producers, regardless of their computational experience, accessibility, or motivation to incorporate AI, has the potential to not only enhance accuracy but also save time and improve results in the long run. Vegetative indices are more informative than raw bands in terms of signal-to-noise ratio and useful for detecting phenotypes in bands of light that are not present in the RGB spectrum (Danilevicz et al. 2021). Incorporating vegetative indices and multispectral imagery should also be considered in future studies.

The correlation analysis revealed a strong relationship between days to anthesis (DTA) and days to tasseling (DTT), with high R^2^ (0.71) and good correspondence across the entire dataset. Repeatability between traits in the same dataset is a more objective metric since it evaluates consistency of an observation that is not due to chance, and is not dependent on another manual measure, which may have its own error, like R^2^. While the combined analysis demonstrated similar moderate repeatability values for DTA (0.651) and DTT (0.541), the repeatability in individual environments varied substantially (0.365 – 0.885) for DTA and (0.141 – 0.790) for DTT. There are two primary potential causes of reducing repeatability. One being measurement errors from humans (DTA) or the CNN model (DTT), the other is a reduced window of genetic variation. DTT has two additional potential causes of reduced repeatability as measured here, the temporal granularity of the flights and the use of classification. Combined DTT could not be revisited or projected forward or backwards like manually estimated DTA can, compressing the variation. For example, in the combined analysis there is a range of 58 to 81 days for DTT, but 52 to 82 days for DTA.

The phenotyping of DTA in Wisconsin was better such that higher repeatabilities in WI were observed despite a smaller range of flowering time (20 days) compared to TX (30 days). The measurement error was especially notable in TXH3 DTT which had both compressed variation in flowering (10 days) due to heat, and lower resolution images (repeatability = 0.141).

When examining the repeatability of DTA, low values were obtained in TX environments (0.141 - 0.549) compared with WI environments (0.365 – 0.885), suggesting substantial measurement error in observations of this trait. This discrepancy may indicate high amounts of error rather than a lack of a strong genetic component to these traits. Repeatability estimates for both DTA (.885) and DTT (.790) were highest in WIH2 and the repeatability of DTT in WIH1 (.572) exceeded that of DTA (.365). Flights in these environments were more frequent during the flowering window and images were taken at a higher resolution. The overall low repeatability estimates for DTA measurements could be caused by human error through discrepancy in visual detection of small pollen grains or subjective determination, while error in DTT was likely from orthomosaic stitching, low image resolution, and lack of flights concurrent with tassel initiation. The relatively high percentage of variance attributed to the Year component for DTT in Texas is likely due to the large variation in ground sampling distance (6.8mm per pixel in 2020 versus 10.9mm per pixel in 2021). Given these findings, future studies should prioritize the acquisition of high-resolution images with near daily flights within the anticipated flowering window of each treatment to enhance data accuracy and repeatability. Furthermore, if high enough resolution was obtained, it is conceivable that anthers could be detected on the tassel, unifying the CNN approach with conventional DTA measures.

This study highlights the potential for temporal phenotyping by extracting additional phenotypic traits throughout the growing season. This could happen in near-real time, only limited by data processing, or as here, retrospectively. This can provide new information and phenotypes on specific plots and genotypes. Notably these new measures can help breeders and plant biologists gain valuable insights into the dynamics of plant development and potentially stress responses when these phenotypes are observable in RGB imagery. Additionally, this approach, once validated by other researchers, may offer opportunities for timely intervention and management decisions for producers; the most obvious example being detasseling decisions for hybrid seed production.

As compared to existing methods of extracting flowering time, establishing a ground truth in a field by counting tassels requires manual counting, which is both labor-intensive and expensive. This study demonstrated that a qualitative visual trait phenotyped in one environment can easily be extended to another using deep learning to reduce future costs and labor. In fact, the improved estimation of Wisconsin, based on Texas-trained data demonstrated that image and data quality can improve estimation accuracy beyond the original training dataset. Phenotyping using UAS imagery may incur initial expenses associated with equipment purchases but will allow for high-throughput phenotyping at a reduced hourly cost. Furthermore, historical imagery allows retrospective data analysis for new phenotypes as they are developed. There is inherent value to a high volume of data, regardless of quality beyond a certain baseline (Lane and Murray 2021). Eventually, as more breeding programs acquire and routinely collect UAS technology, easy-to-use protocols for processing the vast volumes of data produced will be essential in incorporating phenomic data into multi-omic analyses (Chen et al. 2022). It should also be possible to apply this methodology to unoccupied ground systems or manually acquired imagery provided the input images appear similar to our dataset. Automation of tassel detection not only enhances the scalability and speed of phenotyping but also reduces human error and variability. By establishing a visually scored dataset for model training and then validating this trained model using data from unknown genotypes and environments, the challenges associated with accurately detecting the complex flowering phenotypes were addressed.

## Abbreviations

AI: Artificial Intelligence;
CNN: Convolutional Neural Network;
DL: Deep Learning;
DAP: Days after planting;
DTA: Days to anthesis;
DTT: Days to tassel detection;
G2F: Genomes to Fields Initiative;
RCBD: Randomized Complete Block Design;
RGB: Red/Green/Blue;
TXH1: College Station, Texas field 1;
TXH2: College Station, Texas field 2;
TXH3: College Station, Texas field 3;
UAS: Unoccupied aerial system;
UAV: Unoccupied aerial vehicle;
WIH1: Madison, Wisconsin field;
WIH2: Arlington, Wisconsin field;

## Acknowledgments

The authors would like to thank Texas A&M AgriLife Research, USDA, Texas Corn Producers Board, and Iowa Corn Promotion Board. We are also grateful for the support and assistance of all personnel that provided agronomic and technical support under the Texas A&M Quantitative Genetics and Maize Breeding Program. Portions of this research were conducted with the advanced computing resources provided by Texas A&M High Performance Research Computing. ChatGPT-3.5 and ChatGPT-4 were used in the iterative proposal and troubleshooting of written code.

## Conflict of Interest

The authors declare no conflict of interest.

## Supplemental Material

Python .py code and data .xlsx are listed under Supplemental Material. Zipped image files are available upon request.

## Notes

### Competing Interest Statement

The authors have declared no competing interest.

## References

A, S., & Sangeetha, J. (2021). Smart Irrigation and precision farming of paddy field using unmanned ground vehicle and internet of things system. International Journal of Advanced Computer Science and Applications, 12(12). 10.14569/ijacsa.2021.0121254

Aasen, H., Kirchgessner, N., Walter, A., & Liebisch, F. (2020). PhenoCams for field phenotyping: Using very high temporal resolution digital repeated photography to investigate interactions of growth, phenology, and harvest traits. Frontiers in Plant Science, 11. 10.3389/fpls.2020.00593

Adak, A., Murray, S. C., Anderson, S. L., Popescu, S. C., Malambo, L., Romay, M. C., & de Leon, N. (2021). Unoccupied aerial systems discovered overlooked loci capturing the variation of entire growing period in maize. The Plant Genome, 14(2). 10.1002/tpg2.20102

Adak, A., Murray, S. C., Božinović, S., Lindsey, R., Nakasagga, S., Chatterjee, S., Anderson, S. L., & Wilde, S. (2021). Temporal vegetation indices and plant height from remotely sensed imagery can predict grain yield and flowering time breeding value in maize via machine learning regression. Remote Sensing, 13(11), 2141. 10.3390/rs13112141

Akiba, T., Sano, S., Yanase, T., Ohta, T., & Koyama, M. (2019). Optuna. Proceedings of the 25th ACM SIGKDD International Conference on Knowledge Discovery & Data Mining. 10.1145/3292500.3330701

Alzadjali, A., Alali, M. H., Veeranampalayam Sivakumar, A. N., Deogun, J. S., Scott, S., Schnable, J. C., & Shi, Y. (2021). Maize tassel detection from UAV imagery using Deep Learning. Frontiers in Robotics and AI, 8. 10.3389/frobt.2021.600410

Andersen, J. R., Schrag, T., Melchinger, A. E., Zein, I., & Lübberstedt, T. (2005). Validation of DWARF8 polymorphisms associated with flowering time in elite European inbred lines of maize (Zea mays L.). Theoretical and Applied Genetics, 111(2), 206–217. 10.1007/s00122-005-1996-6

Anderson, S. L., Murray, S. C., Malambo, L., Ratcliff, C., Popescu, S., Cope, D., Chang, A., Jung, J., & Thomasson, J. A. (2019). Prediction of maize grain yield before maturity using improved temporal height estimates of unmanned aerial systems. The Plant Phenome Journal, 2(1), 1–15. 10.2135/tppj2019.02.0004

Araus, J. L., & Cairns, J. E. (2014). Field high-throughput phenotyping: The New Crop Breeding Frontier. Trends in Plant Science, 19(1), 52–61. 10.1016/j.tplants.2013.09.008

Barzin, R., Pathak, R., Lotfi, H., Varco, J., & Bora, G. C. (2020). Use of UAS multispectral imagery at different physiological stages for yield prediction and input resource optimization in corn. Remote Sensing, 12(15), 2392. 10.3390/rs12152392

Baum, M. E., Archontoulis, S. V., & Licht, M. A. (2019). Planting date, hybrid maturity, and weather effects on maize yield and crop stage. Agronomy Journal, 111(1), 303–313. 10.2134/agronj2018.04.0297

Buckler, E. S., Holland, J. B., Bradbury, P. J., Acharya, C. B., Brown, P. J., Browne, C., Ersoz, E., Flint-Garcia, S., Garcia, A., Glaubitz, J. C., Goodman, M. M., Harjes, C., Guill, K., Kroon, D. E., Larsson, S., Lepak, N. K., Li, H., Mitchell, S. E., Pressoir, G., … McMullen, M. D. (2009). The genetic architecture of maize flowering time. Science, 325(5941), 714– 718. 10.1126/science.1174276

Chen, C. J., Rutkoski, J., Schnable, J. C., Murray, S. C., Wang, L., Jin, X., Stich, B., Crossa, J., Hayes, B. J., & Zhang, Z. (2022). Role of the genomics–phenomics–agronomy paradigm in plant breeding. Plant Breeding Reviews, 627–673. 10.1002/9781119874157.ch10

Chivasa, W., Mutanga, O., & Burgueño, J. (2021). UAV-based high-throughput phenotyping to increase prediction and selection accuracy in maize varieties under artificial MSV inoculation. Computers and Electronics in Agriculture, 184, 106128. 10.1016/j.compag.2021.106128

Cárcova, J., Uribelarrea, M., Borrás, L., Otegui, M. E., & Westgate, M. E. (2000). Synchronous pollination within and between ears improves kernel set in maize. Crop Science, 40(4), 1056–1061. 10.2135/cropsci2000.4041056x

Danilevicz, M. F., Bayer, P. E., Boussaid, F., Bennamoun, M., & Edwards, D. (2021). Maize yield prediction at an early developmental stage using multispectral images and genotype data for preliminary hybrid selection. Remote Sensing, 13(19), 3976. 10.3390/rs13193976

Daynard, T. B., Tanner, J. W., & Duncan, W. G. (1971). Duration of the grain filling period and its relation to grain yield in corn, zea mays l. Crop Science, 11(1), 45–48. 10.2135/cropsci1971.0011183x001100010015x

Delseny, M., Han, B., & Hsing, Y. I. (2010). High throughput DNA sequencing: The new sequencing revolution. Plant Science, 179(5), 407–422. 10.1016/j.plantsci.2010.07.019

Fitria Widiawati, I., Nugrahapraja, H., & Fajriyah, R. (2018). K-Nearest Neighbor (KNN) analysis on genes expression datasets of maize nested association mapping (NAM) showed confident classification on organ-specific expression. 2018 1st International Conference on Bioinformatics, Biotechnology, and Biomedical Engineering - Bioinformatics and Biomedical Engineering. 10.1109/biomic.2018.8610577

Furbank, R. T., & Tester, M. (2011). Phenomics – technologies to relieve the phenotyping bottleneck. Trends in Plant Science, 16(12), 635–644. 10.1016/j.tplants.2011.09.005

Gano, B., Bhadra, S., Vilbig, J. M., Ahmed, N., Sagan, V., & Shakoor, N. (2024). Drone based imaging sensors, techniques, and applications in plant phenotyping for crop breeding: A comprehensive review. The Plant Phenome Journal, 7(1). 10.1002/ppj2.20100

Izzam, A. (2017). Genetic variability and correlation studies for morphological and yield traits in maize (Zea mays L.). Pure and Applied Biology, 6(4). 10.19045/bspab.2017.600131

Karami, A., Quijano, K., & Crawford, M. (2021). Advancing Tassel Detection and counting: Annotation and algorithms. Remote Sensing, 13(15), 2881. 10.3390/rs13152881

Khan, S. U., Zheng, Y., Chachar, Z., Zhang, X., Zhou, G., Zong, N., Leng, P., & Zhao, J. (2022). Dissection of maize drought tolerance at the flowering stage using genome-wide association studies. Genes, 13(4), 564. 10.3390/genes13040564

Kumar, A., Desai, S. V., Balasubramanian, V. N., Rajalakshmi, P., Guo, W., Balaji Naik, B., Balram, M., & Desai, U. B. (2021). Efficient maize tassel-detection method using UAV based remote sensing. Remote Sensing Applications: Society and Environment, 23, 100549. 10.1016/j.rsase.2021.100549

Kumar, C., Mubvumba, P., Huang, Y., Dhillon, J., & Reddy, K. (2023). Multi-stage corn yield prediction using high-resolution UAV multispectral data and Machine Learning Models. Agronomy, 13(5), 1277. 10.3390/agronomy13051277

Kurtulmuş, F., & Kavdir, İ. (2014). Detecting corn tassels using computer vision and support Vector Machines. Expert Systems with Applications, 41(16), 7390–7397. 10.1016/j.eswa.2014.06.013

Lachowiec, J., Feldman, M. J., Matias, F. I., Lebauer, D., & Gregory, A. (2024). Unoccupied Aerial Systems Adoption in Agricultural Research. 10.22541/essoar.170654380.06877585/v1

Lane, H. M., & Murray, S. C. (2021). High throughput can produce better decisions than high accuracy when phenotyping plant populations. Crop Science, 61(5), 3301–3313. 10.1002/csc2.20514

LeCun, Y., Bengio, Y., & Hinton, G. (2015). Deep learning. Nature, 521(7553), 436–444. 10.1038/nature14539

Lima, D. C., Aviles, A. C., Alpers, R. T., Perkins, A., Schoemaker, D. L., Costa, M., Michel, K. J., Kaeppler, S., Ertl, D., Romay, M. C., Gage, J. L., Holland, J., Beissinger, T., Bohn, M., Buckler, E., Edwards, J., Flint-Garcia, S., Gore, M. A., Hirsch, C. N., … de Leon, N. (2023). 2020-2021 field seasons of Maize GxE project within the genomes to fields initiative. BMC Research Notes, 16(1). 10.1186/s13104-023-06430-y

Liu, Y., Cen, C., Che, Y., Ke, R., Ma, Y., & Ma, Y. (2020). Detection of maize tassels from UAV RGB imagery with faster R-CNN. Remote Sensing, 12(2), 338. 10.3390/rs12020338

Lu, H., & Cao, Z. (2020). TASSELNETV2+: A fast implementation for high-throughput plant counting from high-resolution RGB imagery. Frontiers in Plant Science, 11. 10.3389/fpls.2020.541960

Lu, H., Cao, Z., Xiao, Y., Zhuang, B., & Shen, C. (2017). TasselNet: Counting maize tassels in the wild via local counts regression network. Plant Methods, 13(1). 10.1186/s13007-017-0224-0

Mace, E. S., Hunt, C. H., & Jordan, D. R. (2013). Supermodels: Sorghum and maize provide mutual insight into the genetics of flowering time. Theoretical and Applied Genetics, 126(5), 1377–1395. 10.1007/s00122-013-2059-z

Matias, F. I., Caraza Harter, M. V., & Endelman, J. B. (2020). Fieldimager: An R package to analyze orthomosaic images from agricultural field trials. The Plant Phenome Journal, 3(1). 10.1002/ppj2.20005

Minervini, M., Scharr, H., & Tsaftaris, S. A. (2015). Image analysis: The new bottleneck in plant phenotyping [applications corner]. IEEE Signal Processing Magazine, 32(4), 126–131. 10.1109/msp.2015.2405111

Mirnezami, S. V., Srinivasan, S., Zhou, Y., Schnable, P. S., & Ganapathysubramanian, B. (2021). Detection of the progression of anthesis in field-grown maize tassels: A case study. Plant Phenomics, 2021. 10.34133/2021/4238701

Murcia, H. F., Tilaguy, S., & Ouazaa, S. (2021). Development of a low-cost system for 3D Orchard Mapping Integrating UGV and Lidar. Plants, 10(12), 2804. 10.3390/plants10122804

Poland, J. A., & Rife, T. W. (2012). Genotyping by sequencing for Plant Breeding and Genetics. The Plant Genome, 5(3). 10.3835/plantgenome2012.05.0005

Pugh, N. A., Horne, D. W., Murray, S. C., Carvalho, G., Malambo, L., Jung, J., Chang, A., Maeda, M., Popescu, S., Chu, T., Starek, M. J., Brewer, M. J., Richardson, G., & Rooney, W. L. (2018). Temporal estimates of crop growth in sorghum and maize breeding enabled by unmanned aerial systems. The Plant Phenome Journal, 1(1), 1–10. 10.2135/tppj2017.08.0006

Rattalino Edreira, J. I., Budakli Carpici, E., Sammarro, D., & Otegui, M. E. (2011). Heat stress effects around flowering on kernel set of temperate and tropical maize hybrids. Field Crops Research, 123(2), 62–73. 10.1016/j.fcr.2011.04.015

Ren, S., He, K., Girshick, R., & Sun, J. (2015). Faster R-CNN: Towards real-time object detection with region proposal networks. IEEE Transactions on Pattern Analysis and Machine Intelligence, 39(6), 1137–1149. 10.1109/tpami.2016.2577031

Rincent, R., Charpentier, J.-P., Faivre-Rampant, P., Paux, E., Le Gouis, J., Bastien, C., & Segura, V. (2018). Phenomic selection is a low-cost and high-throughput method based on indirect predictions: Proof of concept on wheat and poplar. G3 Genes|Genomes|Genetics, 8(12), 3961–3972. 10.1534/g3.118.200760

Rodene, E., Fernando, G. D., Piyush, V., Ge, Y., Schnable, J. C., Ghosh, S., & Yang, J. (2024). Image filtering to improve maize tassel detection accuracy using machine learning algorithms. Sensors, 24(7), 2172. 10.3390/s24072172

Shao, M., Nie, C., Cheng, M., Yu, X., Bai, Y., Ming, B., Song, H., & Jin, X. (2021). Quantifying effect of tassels on near-ground maize canopy RGB images using deep learning segmentation algorithm. Precision Agriculture, 23(2), 400–418. 10.1007/s11119-021-09842-7

Shao, M., Nie, C., Zhang, A., Shi, L., Zha, Y., Xu, H., Yang, H., Yu, X., Bai, Y., Liu, S., Cheng, M., Lin, T., Cui, N., Wu, W., & Jin, X. (2023). Quantifying effect of maize tassels on LAI estimation based on multispectral imagery and machine learning methods. Computers and Electronics in Agriculture, 211, 108029. 10.1016/j.compag.2023.108029

Song, P., Wang, J., Guo, X., Yang, W., & Zhao, C. (2021). High-throughput phenotyping: Breaking through the bottleneck in future crop breeding. The Crop Journal, 9(3), 633–645. 10.1016/j.cj.2021.03.015

Sunoj, S., Cho, J., Guinness, J., van Aardt, J., Czymmek, K. J., & Ketterings, Q. M. (2021). Corn grain yield prediction and mapping from unmanned aerial system (UAS) multispectral imagery. Remote Sensing, 13(19), 3948. 10.3390/rs13193948

Tirado, S. B., Hirsch, C. N., & Springer, N. M. (2020). UAV based imaging platform for monitoring maize growth throughout development. Plant Direct, 4(6). 10.1002/pld3.230

Ubbens, J. R., & Stavness, I. (2017). Deep Plant Phenomics: A deep learning platform for complex plant phenotyping tasks. Frontiers in Plant Science, 8. 10.3389/fpls.2017.01190

Warrington, I. J., & Kanemasu, E. T. (1983). Corn growth response to temperature and photoperiod I. Seedling emergence, tassel initiation, and anthesis1. Agronomy Journal, 75(5), 749–754. 10.2134/agronj1983.00021962007500050008x

Worku, M., Makumbi, D., Beyene, Y., Das, B., Mugo, S., Pixley, K., Bänziger, M., Owino, F., Olsen, M., Asea, G., & Prasanna, B. M. (2016). Grain yield performance and flowering synchrony of Cimmyt’s tropical maize (Zea mays L.) parental inbred lines and single crosses. Euphytica, 211(3), 395–409. 10.1007/s10681-016-1758-3

Wu, G., Miller, N. D., de Leon, N., Kaeppler, S. M., & Spalding, E. P. (2019). Predicting Zea mays flowering time, yield, and kernel dimensions by analyzing aerial images. Frontiers in Plant Science, 10. 10.3389/fpls.2019.01251

Yu, X., Yin, D., Nie, C., Ming, B., Xu, H., Liu, Y., Bai, Y., Shao, M., Cheng, M., Liu, Y., Liu, S., Wang, Z., Wang, S., Shi, L., & Jin, X. (2022). Maize tassel area dynamic monitoring based on near-ground and UAV RGB images by U-Net model. Computers and Electronics in Agriculture, 203, 107477. 10.1016/j.compag.2022.107477

Zan, X., Zhang, X., Xing, Z., Liu, W., Zhang, X., Su, W., Liu, Z., Zhao, Y., & Li, S. (2020). Automatic detection of maize tassels from UAV images by combining random forest classifier and VGG16. Remote Sensing, 12(18), 3049. 10.3390/rs12183049

Zhang, W., Wu, S., Wen, W., Lu, X., Wang, C., Gou, W., Li, Y., Guo, X., & Zhao, C. (2023). Three-dimensional branch segmentation and phenotype extraction of Maize Tassel based on Deep Learning. Plant Methods, 19(1). 10.1186/s13007-023-01051-9

Zhu, X., Leiser, W. L., Hahn, V., & Würschum, T. (2021). Phenomic selection is competitive with genomic selection for breeding of complex traits. The Plant Phenome Journal, 4(1). 10.1002/ppj2.20027

