## Supplementary Information for "Deep Learning-Based High-Throughput Phenotyping Of Maize (*Zea mays* L.) Tasseling From Uas Imagery Across Environments"

**Supplementary Table 1. ANOVA results for DTA of the TXH1 dataset**

|  | vcov | sdcor | Percent | RMSE | Repeatability | RSquared | Trait |
| --- | --- | --- | --- | --- | --- | --- | --- |
| Pedigree | 1.09 | 1.05 | .075 |  |  |  |  |
| Row | .017 | .131 | .001 |  |  |  |  |
| Range | .1 | .316 | .007 |  |  |  |  |
| Rep | <.001 | <.001 | <.001 |  |  |  |  |
| Year | 11.7 | 3.42 | .795 |  |  |  |  |
| Residual | 1.80 | 1.34 | .122 | 1.203 | .549 | .878 | TXH1 DTA |

**Supplementary Table 2. ANOVA results for DTT of the TXH1 dataset**

|  | vcov | sdcor | Percent | RMSE | Repeatability | RSquared | Trait |
| --- | --- | --- | --- | --- | --- | --- | --- |
| Pedigree | .61 | .781 | .059 |  |  |  |  |
| Row | <.001 | <.001 | <.001 |  |  |  |  |
| Range | .482 | .694 | .047 |  |  |  |  |
| Rep | .192 | .439 | .018 |  |  |  |  |
| Year | 5.60 | 2.37 | .541 |  |  |  |  |
| Residual | 5.05 | 2.25 | .366 | 1.748 | .261 | .665 | TXH1 DTT |

**Supplementary Table 3. ANOVA results for DTA of the TXH2 dataset**

|  | vcov | sdcor | Percent | RMSE | Repeatability | RSquared | Trait |
| --- | --- | --- | --- | --- | --- | --- | --- |
| Pedigree | 1.19 | 1.09 | .091 |  |  |  |  |
| Row | .085 | .292 | .007 |  |  |  |  |
| Range | .224 | .473 | .017 |  |  |  |  |
| Rep | .020 | .140 | .002 |  |  |  |  |
| Year | 9.27 | 3.04 | .71 |  |  |  |  |
| Residual | 2.26 | 1.50 | .174 | 1.34 | .512 | .826 | TXH2 DTA |

**Supplementary Table 4. ANOVA results for DTT of the TXH2 dataset**

|  | vcov | sdcor | Percent | RMSE | Repeatability | RSquared | Trait |
| --- | --- | --- | --- | --- | --- | --- | --- |
| Pedigree | 1.49 | 1.22 | .108 |  |  |  |  |
| Row | .194 | .440 | .014 |  |  |  |  |
| Range | .679 | .824 | .049 |  |  |  |  |
| Rep | .001 | .037 | <.001 |  |  |  |  |
| Year | 6.36 | 2.52 | .462 |  |  |  |  |
| Residual | 5.05 | 2.246 | .366 | 2.04 | .372 | .634 | TXH2 DTT |

**Supplementary Table 5 ANOVA results for DTA of the TXH3 dataset**

|  | vcov | sdcor | Percent | RMSE | Repeatability | RSquared | Trait |
| --- | --- | --- | --- | --- | --- | --- | --- |
| Pedigree | 1.07 | 1.03 | .172 |  |  |  |  |
| Row | .049 | .222 | .008 |  |  |  |  |
| Range | .988 | .994 | .309 |  |  |  |  |
| Rep | .323 | .568 | .052 |  |  |  |  |
| Year | 1.91 | 1.38 | .309 |  |  |  |  |
| Residual | 1.84 | 1.36 | .298 | 1.16 | .536 | .702 | TXH3 DTA |

**Supplementary Table 6. ANOVA results for DTT of the TXH3 dataset**

|  | vcov | sdcor | Percent | RMSE | Repeatability | RSquared | Trait |
| --- | --- | --- | --- | --- | --- | --- | --- |
| Pedigree | .216 | .465 | .025 |  |  |  |  |
| Row | <.001 | <.001 | <.001 |  |  |  |  |
| Range | .134 | .366 | .016 |  |  |  |  |
| Rep | .039 | .200 | .005 |  |  |  |  |
| Year | 5.49 | 2.34 | .646 |  |  |  |  |
| Residual | 2.62 | 1.62 | .308 | 1.55 | .141 | .691 | TXH3 DTT |

**Supplementary Table 7. ANOVA results for DTA of the WIH1 dataset**

|  | vcov | sdcor | Percent | RMSE | Repeatability | RSquared | Trait |
| --- | --- | --- | --- | --- | --- | --- | --- |
| Pedigree | .441 | .664 | .191 |  |  |  |  |
| Row | .276 | .525 | .119 |  |  |  |  |
| Range | <.001 | <.001 | <.001 |  |  |  |  |
| Rep | .056 | .237 | .024 |  |  |  |  |
| Residual | 1.53 | 1.24 | .665 | 1.08 | .365 | .335 | WIH1 DTA |

**Supplementary Table 8. ANOVA results for DTT of the WIH1 dataset**

|  | vcov | sdcor | Percent | RMSE | Repeatability | RSquared | Trait |
| --- | --- | --- | --- | --- | --- | --- | --- |
| Pedigree | .252 | .502 | .370 |  |  |  |  |
| Row | .053 | .231 | .078 |  |  |  |  |
| Range | <.001 | <.001 | <.001 |  |  |  |  |
| Rep | <.001 | <.001 | <.001 |  |  |  |  |
| Residual | .377 | .614 | .552 | .480 | .572 | .448 | WIH1 DTT |

**Supplementary Table 9. ANOVA results for DTA of the WIH2 dataset**

|  | vcov | sdcor | Percent | RMSE | Repeatability | RSquared | Trait |
| --- | --- | --- | --- | --- | --- | --- | --- |
| Pedigree | 1.82 | 1.349 | .749 |  |  |  |  |
| Row | .138 | .372 | .057 |  |  |  |  |
| Range | <.001 | <.001 | <.001 |  |  |  |  |
| Rep | <.001 | .007 | <.001 |  |  |  |  |
| Residual | .472 | .687 | .194 | .374 | .885 | .806 | WIH2 DTA |

**Supplementary Table 10. ANOVA results for DTT of the WIH2 dataset**

|  | vcov | sdcor | Percent | RMSE | Repeatability | RSquared | Trait |
| --- | --- | --- | --- | --- | --- | --- | --- |
| Pedigree | 1.65 | 1.285 | .591 |  |  |  |  |
| Row | .215 | .463 | .077 |  |  |  |  |
| Range | .033 | .182 | .012 |  |  |  |  |
| Rep | .017 | .129 | .006 |  |  |  |  |
| Residual | .876 | .936 | .314 | .589 | .790 | .686 | WIH2 DTT |

**Supplementary Table 11**. Comparisons between traditional and UAS-based phenotyping methods.

|  | **Traditional field-based phenotyping** | **UAS-based high throughput phenotyping** |
| --- | --- | --- |
| **Cost** | Hourly wages for personnel. Specialized personnel are not necessary but likely improve accuracy. | Phantom 4 Pro v2 Drone: $1500  Agisoft Metashape Professional (Education Edition): $549/year  GPS for ground control points: $1000+  Desktop computer with adequate CPU/GPU/RAM/storage: $2000+ |
| **Time** | Scoring and recording data for 200 hybrids requires between 2-2.5 hours of time for one rater (provided field is dry enough for walking). | Each UAS flight takes roughly 30 minutes; mosaicking process requires between 4 to 8 hours to stitch images together (mostly computational, approximately 2 hours of manual input); and manual scoring of each hybrid from a mosaic requires approximately 6 hours per mosaic. |
| **Accuracy** | Rater bias is subject to level of expertise, environmental factors, lighting conditions, weather, and fatigue. | Orthomosaics provide uniform arrangement of plants to be scored; subject to lighting conditions during UAS flight, quality of image stitching, and (to a lesser degree) rater bias. |
| **Resolution** | No image resolution limitations, however limited to lower and mid-level leaves at the end of plot, the interior of plots is more challenging to observe; canopy can be difficult to phenotype when plants are at maximum height. | Pixel size of mosaics is determined by UAS flying height, speed and camera resolution and ability; visible plant area restricted to canopy and some mid-level leaves. |
| **Scale** | Limited by available and trained personnel in the field. | Limited by the available number of batteries for UAS and willingness of operator in the field. Limited by available and trained personnel in the lab. |

**References**

1	DeSalvio, A. J., Adak, A., Murray, S. C., Wilde, S. C., & Isakeit, T. (2022). Phenomic data-facilitated rust and senescence prediction in maize using machine learning algorithms. Scientific reports, 12(1), 7571.
